# Fine-tuning Polygenic Risk Scores with GWAS Summary Statistics

**DOI:** 10.1101/810713

**Authors:** Zijie Zhao, Yanyao Yi, Yuchang Wu, Xiaoyuan Zhong, Yupei Lin, Timothy J. Hohman, Jason Fletcher, Qiongshi Lu

## Abstract

Polygenic risk scores (PRSs) have wide applications in human genetics research. Notably, most PRS models include tuning parameters which improve predictive performance when properly selected. However, existing model-tuning methods require individual-level genetic data as the training dataset or as a validation dataset independent from both training and testing samples. These data rarely exist in practice, creating a significant gap between PRS methodology and applications. Here, we introduce PUMAS (Parameter-tuning Using Marginal Association Statistics), a novel method to fine-tune PRS models using summary statistics from genome-wide association studies (GWASs). Through extensive simulations, external validations, and analysis of 65 traits, we demonstrate that PUMAS can perform a variety of model-tuning procedures (e.g. cross-validation) using GWAS summary statistics and can effectively benchmark and optimize PRS models under diverse genetic architecture. On average, PUMAS improves the predictive R^2^ by 205.6% and 62.5% compared to PRSs with arbitrary p-value cutoffs of 0.01 and 1, respectively. Applied to 211 neuroimaging traits and Alzheimer’s disease, we show that fine-tuned PRSs will significantly improve statistical power in downstream association analysis. We believe our method resolves a fundamental problem without a current solution and will greatly benefit genetic prediction applications.

## Introduction

Accurate prediction of complex traits with genetic data is a major goal in human genetics research and precision medicine.^1^ In the past decade, advancements in genotyping and imputation techniques have greatly accelerated discoveries in genome-wide association studies (GWASs) for numerous complex diseases and traits.^2^ These data have also enabled statistical learning applications that leverage genome-wide data in genetic risk prediction.^3-8^ However, despite these advances, it remains challenging to access, store, and process individual-level genetic data at a large scale due to privacy concerns and high computational burden. With increasingly accessible GWAS summary statistics for a variety of complex traits,^9^ polygenic risk scores (PRSs) that use marginal association statistics as input enjoy great popularity and have had success in diverse applications.^10-12^

With great popularity there also come great challenges. Prediction accuracy of PRS remains moderate for most phenotypes.^13^ Methods have been developed to improve PRS performance by explicitly modeling linkage disequilibrium (LD),^14^ incorporating functional annotations and pleiotropy,^15,16^ and improving effect estimates through statistical shrinkage.^17^ Notably, most PRS models have tuning parameters, including the p-value threshold in traditional PRS, the penalty strength in penalized regression models, and the proportion of causal variants in LDpred.^14^ Tuning parameters are very common in predictive modeling. When properly selected, these parameters add flexibility to the model and improve prediction accuracy. This is a well-understood problem with a rich literature – a well-known solution is cross-validation.^18^ However, most model-tuning methods require individual-level genetic data either as the training dataset or as a validation dataset independent from both the input GWAS and the testing samples. In practice, these data rarely exist, especially when PRS is generated using GWAS summary statistics in the public domain. This has created a significant gap between current conventions in PRS construction and optimal methodologies. Without a method to fine-tune models using summary statistics, it is challenging to benchmark and optimize PRS, thus limiting its clinical utility.

We introduce PUMAS (Parameter-tuning Using Marginal Association Statistics), a novel method to fine-tune PRS models using GWAS summary data. As a general framework, PUMAS can conduct a variety of model-tuning procedures on PRS, including training-testing data split, cross-validation, and repeated learning. Through extensive simulations on realistic genetic architecture, we demonstrate that the performance of PUMAS is as good as methods based on individual-level data. Additionally, we apply PUMAS to GWAS traits with distinct types of genetic architecture and validate our results using well-powered external datasets. Further, we systematically benchmark and optimize PRS for numerous diseases and traits and showcase the immediate benefits of fine-tuned PRSs in downstream applications.

## Results

### Method overview

Here, we outline the PUMAS framework. Detailed derivations and technical discussions are included in the **Methods** section. There are two key steps in our proposed model-tuning framework (**Figure 1**). First, we sample marginal association statistics for a subset of individuals based on the complete GWAS summary data. Using this approach, we can generate summary statistics for independent training and testing sets without actually partitioning the samples. Second, we propose an approach to evaluate the predictive performance (e.g. predictive R^2^) of PRS using summary statistics in the validation set so that we can select the best model based on its superior performance. These two steps together make it possible to select the best-performing model with only one set of GWAS summary statistics as input.

**Figure 1.**
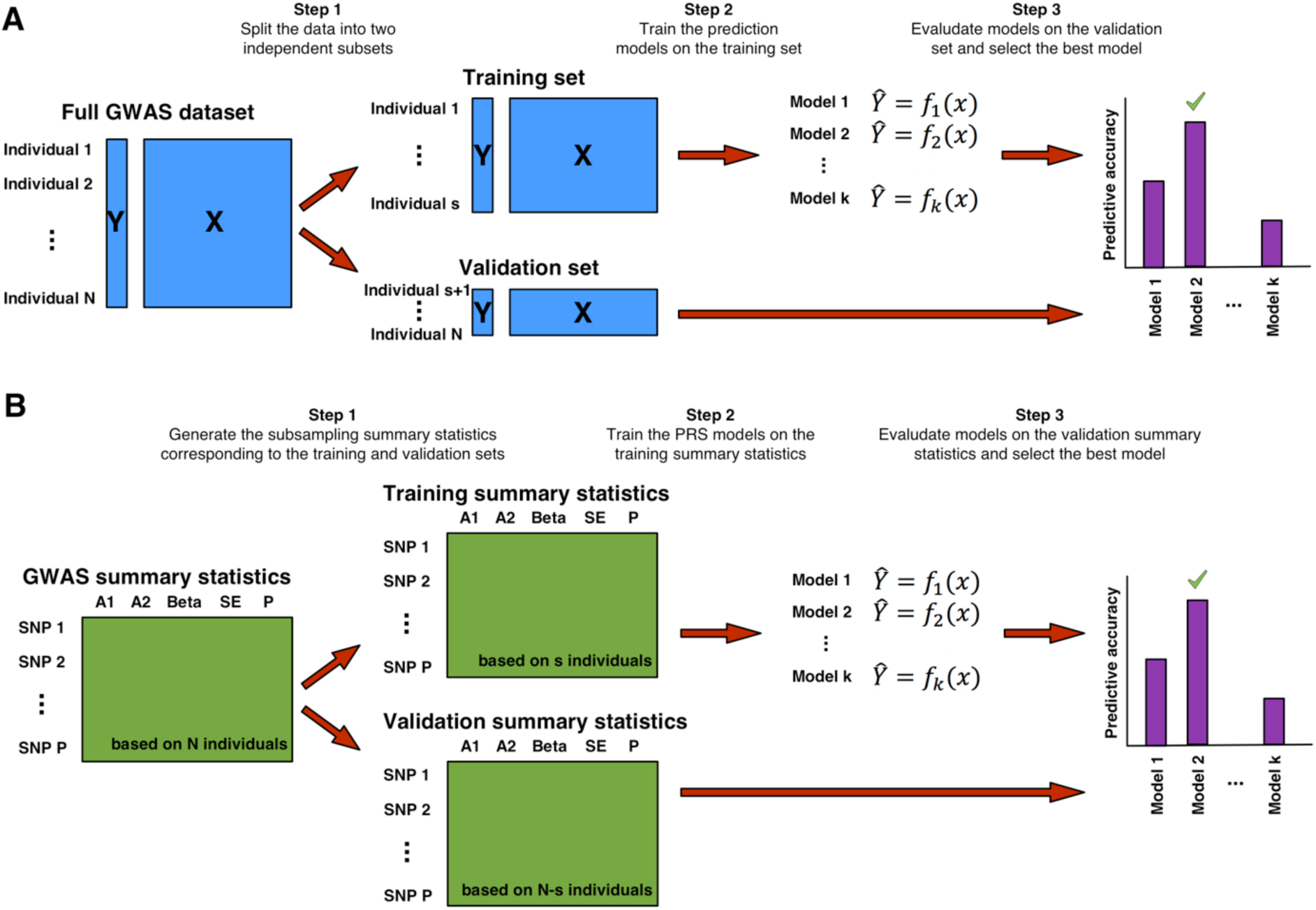
A workflow of model-tuning strategies. **(A)** Traditional approaches split individual-level data into training and validation subsets to fine-tune prediction models. **(B)** Our method directly generates training and testing summary statistics without using individual-level information and use simulated summary statistics as input to select the best model.

### Simulation results

We conducted simulations to investigate if PUMAS can achieve similar performance compared to classic model-tuning procedures. We simulated both genotype and phenotype data with varying sample size, proportion of causal variants, and heritability (**Methods**). We used these data to calculate marginal association statistics and ranked SNPs based on association p-values. Next, we applied PUMAS to perform 4-fold repeated learning on marginal association statistics and selected the optimal number of SNPs to include in the prediction model by maximizing the average *R*^2^ across folds. Additionally, we implemented a traditional repeated learning approach with the same simulated individual-level data as a reference. The two approaches yielded highly consistent results (**Figure 2** and **Supplementary Figure 1**; **Supplementary Table 1**). Since SNP effects were randomly sampled from a normal distribution, it is expected that some weak effects are dropped from the optimal PRS model, leading to a lower number of selected predictors compared to the number of true causal variants, especially when sample size and heritability are low. Across all simulation settings, our summary statistics-based approach showed nearly identical results compared to a state-of-the-art model-tuning approach based on individual-level data and could effectively select the optimal tuning parameter (i.e. number of SNPs in the PRS).

**Figure 2.**
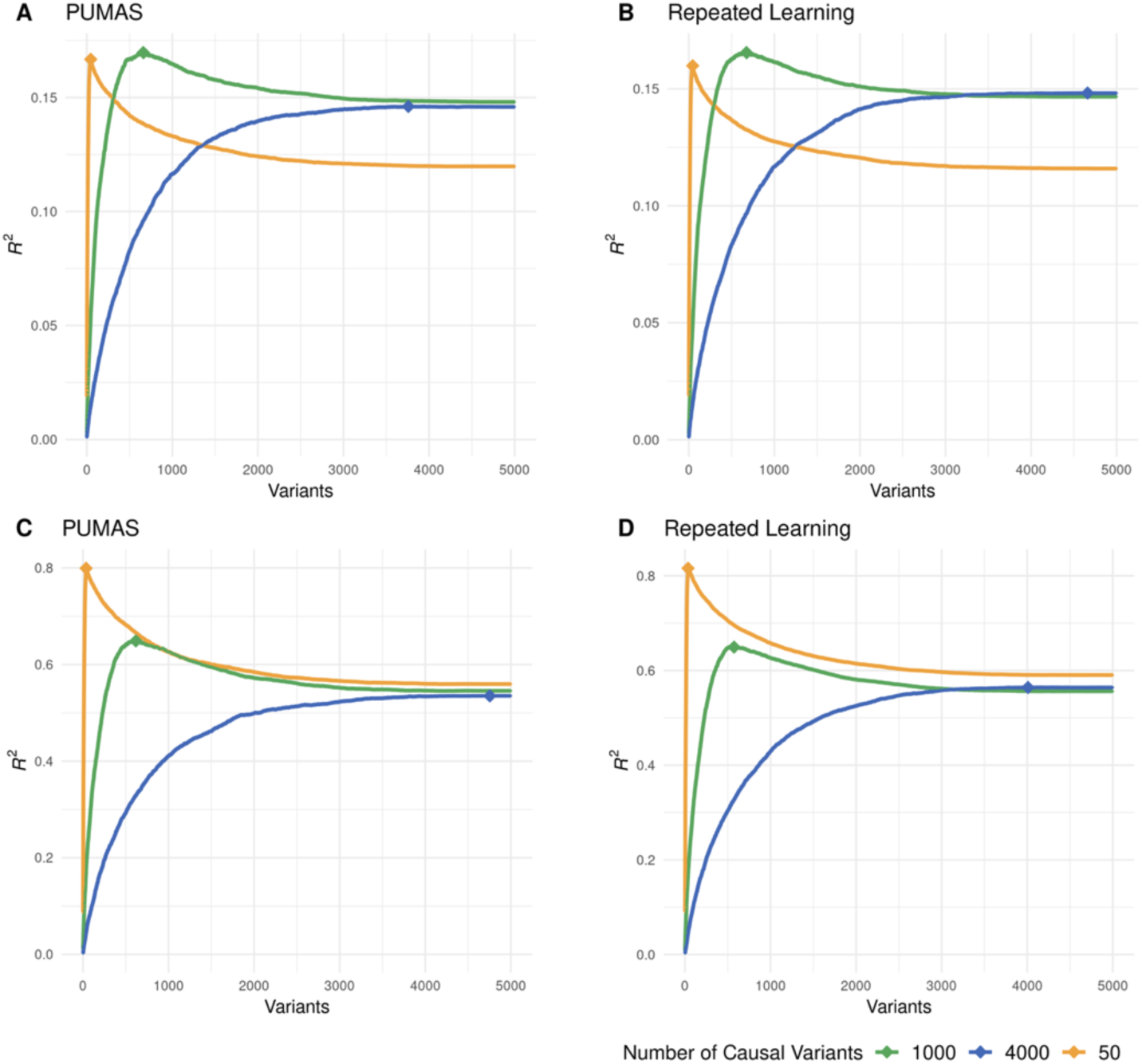
Comparing the performance of two model-tuning strategies. The first two panels illustrate the model-tuning performance of **(A)** PUMAS and **(B)** repeated learning approach based on individual-level data (N=100,000, *h*^*2*^=0.2). Panels **(C-D)** shows the performance of PUMAS and repeated learning on 20,000 samples when heritability is 0.8. The X-axis shows the value of tuning parameter (i.e. number of variants to include in the model) in these simulations. The Y-axis shows the predictive *R*^2^. The three curves in each panel represent three different levels of sparsity and genetic architecture. Results for other settings are summarized in **Supplementary Figure 1**.

### PUMAS effectively fine-tunes PRS models based on genetic architecture

Next, we demonstrate our method’s performance using a gold-standard approach – we apply PUMAS to the summary statistics from well-powered GWASs to select the optimal p-value cutoffs in PRS models and validate their performance on large independent cohorts. First, we applied PUMAS to a recent GWAS of educational attainment (EA) conducted by the Social Science Genetic Association Consortium (N=742,903).^19^ 4,775 samples with European ancestry in the National Longitudinal Study of Adolescent to Adult Health (Add Health)^20^ and 10,214 European samples in the Health and Retirement Study (HRS)^21^ were used as two independent validation sets to assess the predictive performance of EA PRS. We used GWAS of Alzheimer’s disease (AD) as a second example. We applied PUMAS to the stage-1 summary statistics from the 2013 study conducted by the International Genomics of Alzheimer’s Project (IGAP; N=54,162) to optimize PRS models for AD. These PRSs were then evaluated on 7,050 independent samples^22^ from the Alzheimer’s Disease Genetics Consortium (ADGC) and 355,583 samples in the UK Biobank with a family history-based proxy phenotype for AD (**Methods**).^23^

Our summary statistics-based analyses showed highly consistent results compared with external validations (**Figure 3**; **Supplementary Table 2**). Our analysis clearly suggested that a model with a large number of SNPs tend to be more predictive for EA, a pattern validated in both Add Health and HRS cohorts. The EA PRS based on p-value cutoffs of 0.8, 0.8, and 0.7 were the most predictive models suggested by PUMAS, HRS, and Add Health cohorts, respectively. Results on AD were also consistent between PUMAS and external validations. The optimal p-value cutoffs suggested by PUMAS, ADGC validation, and UK Biobank validation were 5e-7, 5e-8, and 1e-10, respectively. PRS models based on p-value cutoffs more stringent than 1e-5 showed good predictive performance in two validation sets for AD. Notably, as more SNPs are included in the model, predictive performance of PRS sharply declines. Our model-tuning results based on GWAS summary statistics accurately predicted this pattern. Additionally, since we used an AD proxy phenotype in the UK Biobank, the reduced predictive R^2^ is expected. But the trend of predictive performance remained consistent with the validation result in case-control data from the ADGC.

**Figure 3.**
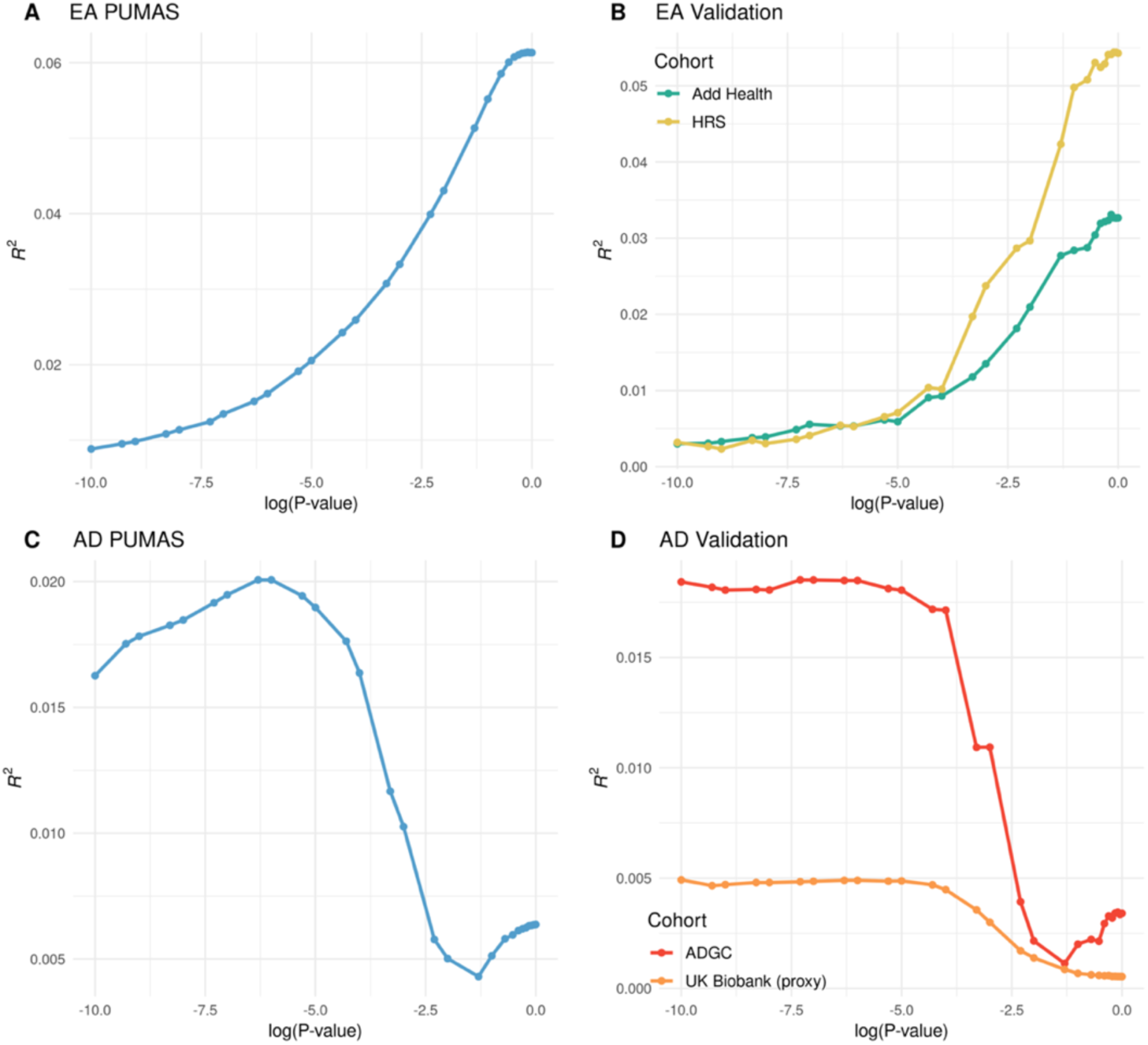
Model-tuning performance on real GWAS data. **(A)** PUMAS performance on the EA training set. **(B)** Prediction performance on two validation sets for EA. **(C)** PUMAS performance on the AD training set. **(D)** Prediction performance on two validation sets for AD. The X-axis shows the log-transformed p-value cutoffs in PRS which is the tuning parameter of interest. The Y-axis indicates predictive *R*^2^. EA: educational attainment; AD: Alzheimer’s disease.

EA is known to be extremely polygenic – more than 1,200 independent genetic associations have been identified for EA to date.^19^ AD has a very different genetic architecture compared to EA. The *APOE* locus has an unusually large effect on AD risk.^24^ In addition to *APOE*, about 30 independent loci have been implicated in AD GWASs.^25^ Our method correctly suggested that the EA PRS would perform better if more SNPs are in the model (87,985 SNPs were included) while a substantially sparser model with 29 SNPs would yield better predictive performance for AD. These results showcased our method’s ability to adaptively choose the optimal tuning parameter for traits with different patterns of genetic architecture. These results also highlighted the importance of model tuning. An AD PRS based on an arbitrary p-value cutoff of 0.01 can have a 5-fold reduction in predictive *R*^2^ compared to the fine-tuned PRS.

### Some technical considerations

We discuss two unique technical issues that may arise in summary statistics-based model tuning. First, sample sizes for different SNPs in a GWAS meta-analysis may vary due to technical differences across cohorts. However, it is not uncommon for a GWAS to only report the maximum sample size. Here, we investigate the robustness of PUMAS when sample size is mis-specified. We use two GWAS datasets that provided accurate sample size for each SNP: summary statistics for low-density lipoprotein (LDL) cholesterol from the Global Lipids Genetics Consortium (GLGC; N=188,577)^26^ and the same EA GWAS summary statistics we have described before. We compared PUMAS results based on four different approaches. The first approach uses the accurate sample size reported in the summary statistics (‘original’); The second approach removes SNPs with sample size below the 30% quantile of its distribution and uses the accurate sample size for remaining SNPs (‘QCed’). The third and fourth approaches apply the maximum or minimum sample size to all SNPs (‘Uniform large/small N’). For the ‘original’ and ‘QCed’ approaches where precise sample size is available for each SNP, we assigned 25% of the minimal N value as the sample size for the validation dataset and used the remaining samples of each SNP in the training subset. Overall, PUMAS results showed consistent patterns under these four scenarios (**Figure 4A)**. Although the R^2^ estimates can inflate or deflate if the sample size is mis-specified, the optimal p-value cutoffs selected by PUMAS remained stable. Thus, PUMAS can still select the best-performing model even if accurate sample size information is unavailable. In practice, performing quality control to remove SNPs with outlier sample size may make the R^2^ estimates most interpretable.

**Figure 4.**
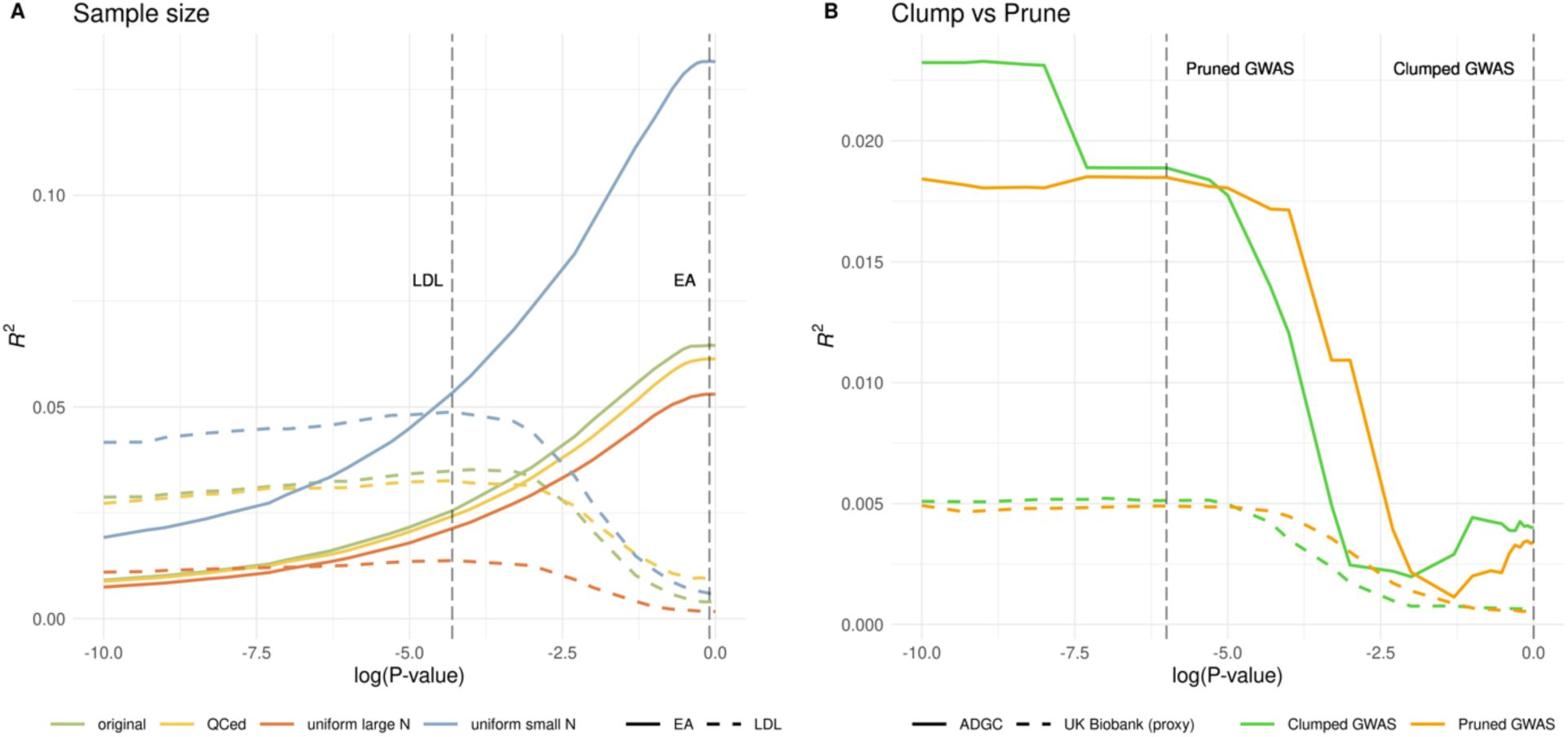
Technical issues involving sample size and LD clumping. **(A)** PUMAS results on LDL cholesterol and EA with various sample size specifications. The two grey dashed lines represent the optimal p-value cutoffs selected by the ‘QCed’ setting for LDL and EA, respectively. **(B)** Predictive performance on external validation for AD PRS based on pruned and clumped summary statistics. Two grey dashed lines mark the optimal p-value cutoffs inferred by PUMAS on pruned and clumped summary statistics. LDL: low-density lipoprotein; EA: educational attainment; AD: Alzheimer’s disease.

The second issue is to see if PUMAS can be applied to clumped GWAS summary statistics. In PRS applications, it is a common practice to clump the data by removing SNPs in strong LD with the most significant SNP in a region. However, since p-values based on the full sample have been used during LD clumping, directly applying the same model-tuning methods to clumped data may lead to information leak and overfitting. We applied PUMAS to clumped summary statistics of the IGAP 2013 AD GWAS (**Supplementary Figure 2**). The model-tuning results in PUMAS were completely inconsistent with the optimal models in external validation (**Figure 4B**), confirming that PUMAS should not be applied to clumped data. However, we note that the predictive curves were very similar in external validations no matter if pruned or clumped data were used as input. Therefore, in practice, it may be plausible to apply PUMAS to pruned GWAS summary data and obtain the optimal p-value threshold. This way, p-values based on the complete sample will not influence the model-tuning procedure. Then, we can apply this selected p-value cutoff with clumped GWAS summary statistics to calculate PRS.

### Benchmarking and optimizing PRS for 65 diseases and traits

Next, we apply PUMAS to provide an atlas of optimized PRSs for complex diseases and traits (**Figure 5**). In total, we analyzed 65 GWASs with available summary statistics and documented each trait’s optimal p-value cutoff and predictive R^2^ (**Supplementary Table 3**). The average gain in predictive R^2^ with our method is 0.0106 (205.6% improvement) and 0.0034 (62.5% improvement) compared to PRSs with p-value cutoffs of 0.01 and 1, respectively (**Supplementary Table 4** and **Supplementary Figure 3**). We annotated the traits into five categories: behavioral/social, metabolic/cardiovascular, psychiatric/neurological, immune, and other. Most behavioral/social traits and psychiatric/neurological disorders had optimal p-value cutoffs between 0.1 and 1 which is consistent with their extreme polygenic genetic architecture. The exceptions include alcoholism (drinks per week), smoking behavior (cigarettes per day), and AD. PRSs with fewer SNPs showed superior performance for these traits. Among immune diseases, systemic lupus erythematosus, primary biliary cirrhosis, rheumatoid arthritis, multiple sclerosis, and eczema all favored a sparse model, while the optimal PRSs for inflammatory bowel diseases and celiac disease had substantially more SNPs. We also note that molecular traits such as blood lipids and 25-hydorxyvitamin D favored sparse PRS models, possibly due to stronger genetic effects and more homogeneous genetic mechanisms. These results also shed light on the differences in the predictive power of diverse types of diseases and traits. PRSs for height, systemic lupus erythematosus, inflammatory bowel diseases, and schizophrenia showed substantially better predictive performance, while the R^2^ for most behavioral/social traits remained moderate despite the large sample size in those studies.

**Figure 5.**
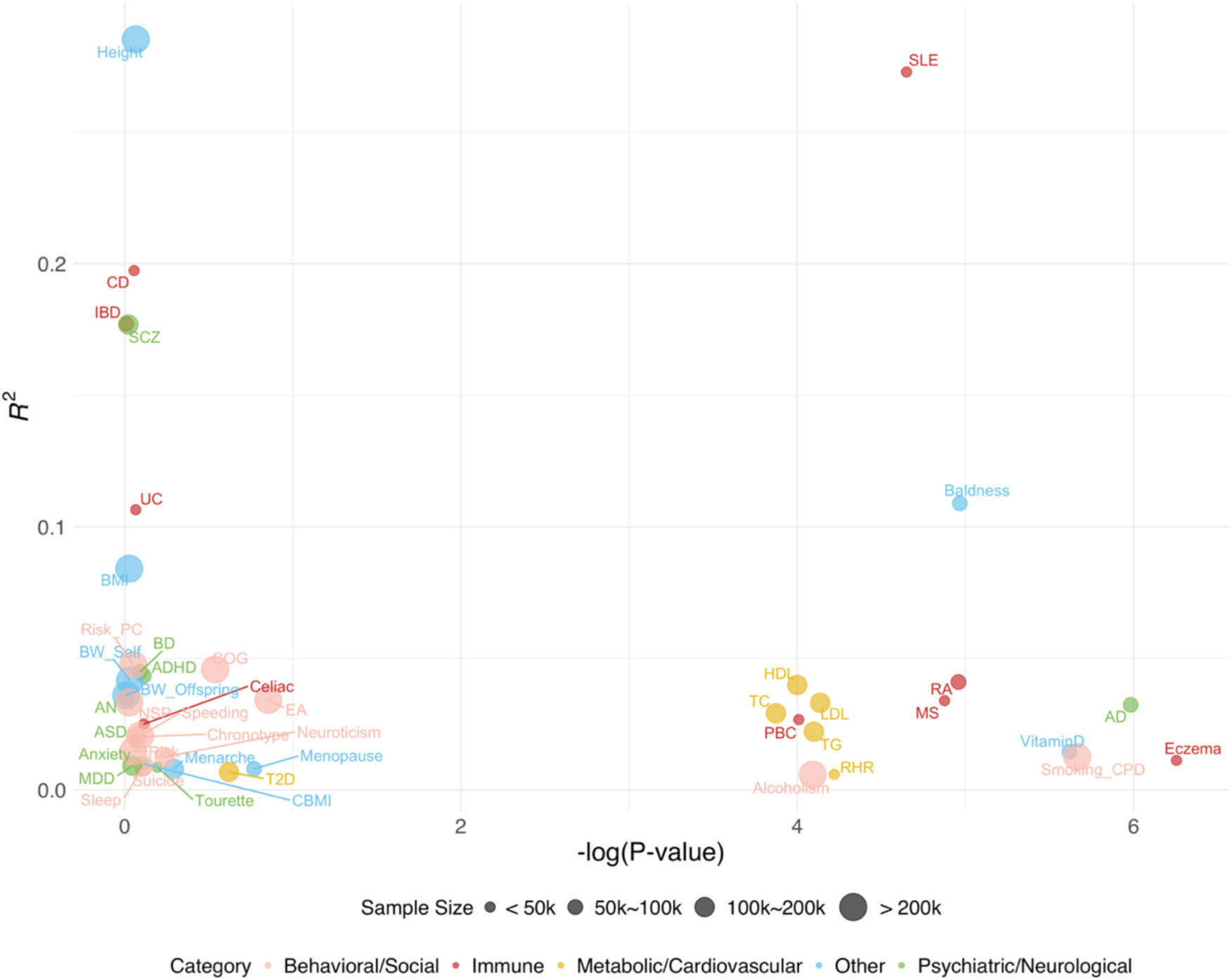
An atlas of optimized PRSs for complex diseases and traits. 45 diseases/traits with optimized R2 > 0.005 are included in the figure. Each circle represents a disease or trait. The size of circles indicates the sample size of the study; colors mark the five trait categories. The X-axis indicates the negative log-transformed p-value cutoff in PRS which is also the tuning parameter of interest. The Y-axis indicates the optimal *R*^2^. Information on all diseases and traits is summarized in **Supplementary Table 3**.

### Identifying neuroimaging associations for AD

Finally, we demonstrate that fine-tuned PRS will lead to power gain in association analysis. We generated PRSs for 211 neuroimaging traits based on two recent studies conducted using samples from the UK Biobank (N=17,706 and 19,629 for diffusion tensor imaging traits and regional volume phenotypes, respectively).^27,28^ We optimized PRS for each imaging trait using PUMAS (**Supplementary Table 5**). For comparison, we also generated PRSs for all traits using an arbitrary p-value cutoff of 0.01. We applied the BADGERS^29^ approach to test associations between 211 neuroimaging trait PRSs with AD in two large, independent AD datasets: the 2019 IGAP GWAS for AD (N=63,926) and the UK Biobank-based GWAS with a proxy phenotype for AD (N=318,773).^23,30^ Samples used in the neuroimaging GWAS were removed from the AD-proxy GWAS to avoid overfitting of PRS models (**Methods**; **Supplementary Table 6** and **Supplementary Figure 4**). Association results in two AD datasets were meta-analyzed to improve statistical power.

The complete association results of 211 neuroimaging traits with AD are summarized in **Supplementary Table 7**. Using fine-tuned PRSs, we identified 2 significant associations with AD under a stringent Bonferroni correction for multiple testing: fornix (cres) / stria terminalis mode of anisotropy (p=1.7E-05) and axial diffusivities (p=2.7E-05) whereby genetic risk for worse white matter integrity in the fornix was associated with risk of AD. No significant associations were identified using PRSs with an arbitrary p-value cutoff (**Figure 6A**). Association p-values based on optimized PRSs were significantly lower than those based on arbitrary PRSs (p=0.03; two-sample Kolmogorov-Smirnov test). Additionally, effect size estimates for top associations were consistent in two independent AD GWASs (**Figure 6B**). Although the effect sizes in two AD studies were not at the same scale due to the difference in AD phenotype definition, effect estimates showed strong concordance between two independent analyses (correlation=0.84). The fornix is a critical white matter tract projecting from the medial temporal lobe where pathology begins in AD, thus it is unsurprising that microstructural changes in the fornix measured with diffusion tensor imaging are observed in mild cognitive impairment and AD.^31-33^ Further, as a negative control, we applied the same analysis to a well-powered breast cancer GWAS (N=228,951).^34^ Results for fine-tuned PRSs and arbitrary PRSs were consistent with the expectation under the null (**Supplementary Figure 5**). No significant associations were identified. These findings demonstrated that our model-tuning approach can increase the statistical power in PRS association analysis.

**Figure 6.**
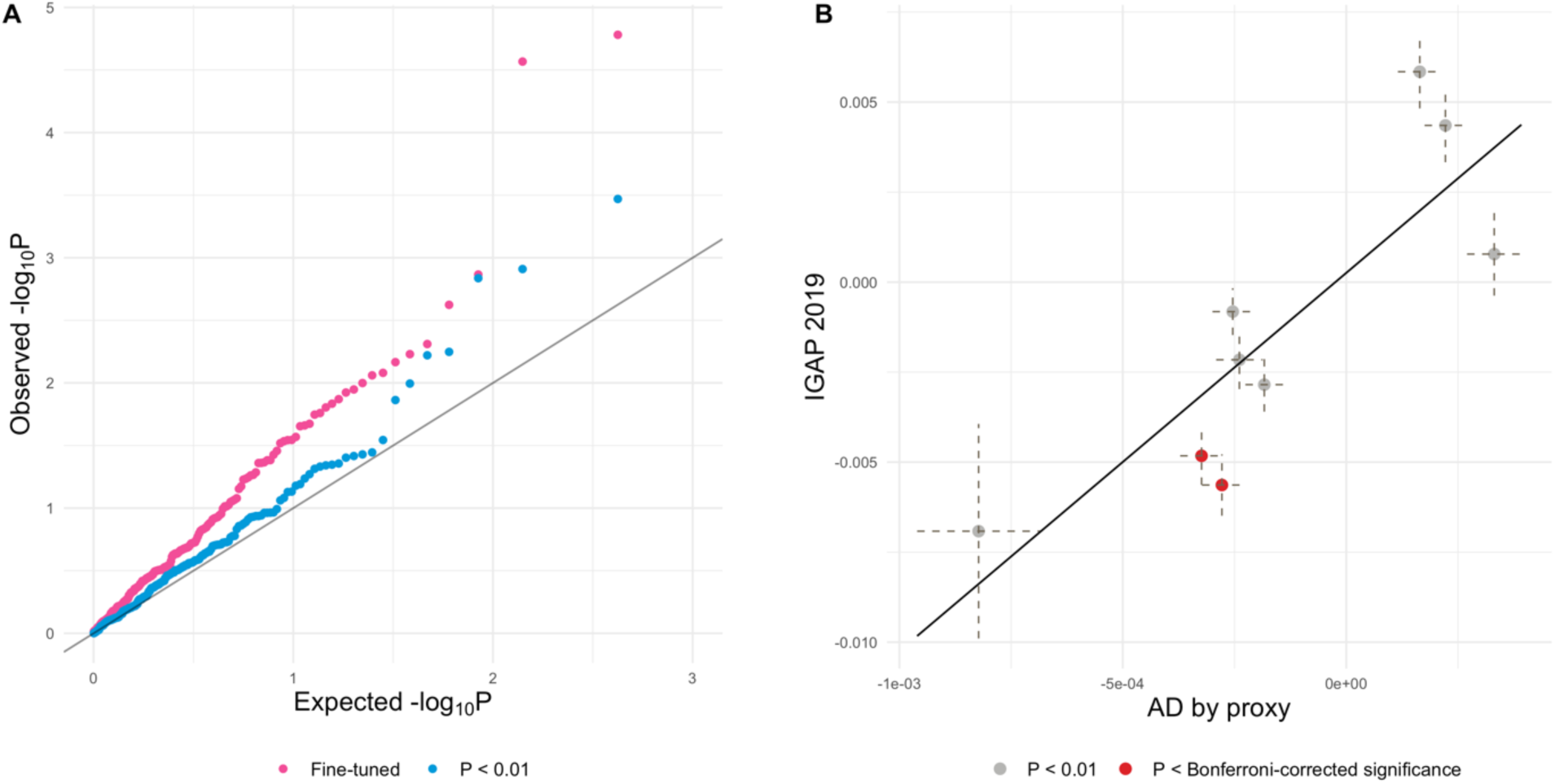
Identifying neuroimaging trait PRSs associated with AD. **(A)** QQ plot for the associations between 211 neuroimaging trait PRSs and AD. P-values were based on the meta-analysis of IGAP 2019 GWAS and the UK Biobank with a proxy AD phenotype. **(B)** Effect size estimates for top associations. Imaging trait PRSs that reached a p-value<0.01 in the meta-analysis are shown in the plot. X-axis: effect sizes of imaging trait PRSs on the AD-proxy phenotype in the UK Biobank; Y-axis: effect sizes on AD in the IGAP 2019 GWAS. Imaging traits whose p-value achieved Bonferroni-corrected significance in the meta-analysis are highlight in red. The dashed lines represent standard error of effect sizes estimates.

## Discussion

Fine-tuning PRS models with GWAS summary statistics has long been considered an impossible task. In this work, we introduced a statistical framework to solve this challenging problem. First, using GWAS summary data as input, PUMAS simulates training and validation summary statistics without accessing individual-level information. Then, PUMAS evaluates and optimizes PRS models on the simulated validation summary statistics. Both steps in the PUMAS framework are statistically rigorous, computationally efficient, and highly novel. Through simulations and analysis of real GWAS data with diverse genetic architecture, we demonstrated that PUMAS can effectively conduct sophisticated model-tuning tasks using GWAS summary statistics. We also showed that optimizing PRSs improves the statistical power in downstream association analysis and identified neuroimaging traits significantly associated with AD.

This work will bring multiple advances to the field. First, it is no longer necessary to leave one dataset out in the GWAS for model tuning purpose. With PUMAS, researchers can safely use effect size estimates from the largest available GWAS for PRS model training, which will lead to improved prediction accuracy. Second, when an independent validation set is not available, most studies in the literature select tuning parameters using one of the two strategies. Some studies fine-tune PRSs on testing samples that are used again in downstream applications, creating an overfitting problem, while other studies use a subset of testing samples to tune the model, reducing the sample size and power in the testing data. PUMAS allows researchers to apply fine-tuned PRS models to the full testing samples, thus avoiding overfitting and improving statistical power. Third, selecting the optimal tuning parameter is not the only application of PUMAS. Given a PRS model, our method allows researchers to calculate cross-validated predictive accuracy, providing a systematic approach to benchmark model performance without requiring external samples.

So far, our analyses have focused on a classic PRS model with pruned SNPs and a varying p-value cutoff that needs to be tuned. Despite the simplicity, it remains one of the most widely used PRS models in the field. However, more sophisticated PRS methods have emerged. Future work will focus on generalizing PUMAS to fine-tune parameters in other PRS models such as LDpred and benchmarking the performance of all models for different traits. Our results have provided strong evidence that it is possible to fine-tune PRS models with GWAS summary data. This new approach, in conjunction with widely available GWAS summary statistics, will have a long-lasting impact on future PRS model development and genetic prediction applications.

## Methods

### Subsampling GWAS association statistics

We assume the quantitative trait Y follows a linear model:

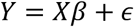

where *X* denotes the SNP genotypes; *β* is a *p*-dimensional vector representing SNP effect sizes; *ϵ* is the error term following a normal distribution with zero mean. Let *y* and *x* = (*x*_1_, …, *x*_*p*_) denote the phenotypic and genotypic data of *N* independent individuals. For simplicity, we assume *y* and *x*_*j*_’s are centered. The summary association statistics in GWAS are obtained from the marginal linear regressions. Then, for *j* = 1, …, *p*, we can denote the regression coefficients and their standard errors as follows

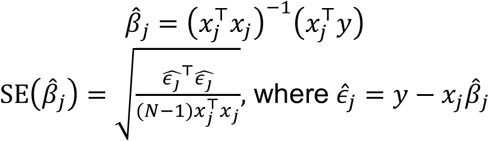

If we have access to the full data set, most model-tuning approaches involve randomly sampling a subset of *N* − *n* individuals as the training set, i.e. *y*^(*tr*)^ and *x*^(*tr*)^. Naturally, the remaining subset of *n* individuals will be the validation dataset denoted as *y*^(*v*)^ and *x*^(*v*)^. When only the summary statistics file based on the full dataset is provided, the traditional model-tuning approaches cannot be implemented. Instead, we propose a method to generate marginal summary statistics for the training and validating datasets from summary statistics of the full dataset. By central limit theorem, as sample size *N* → ∞, we have

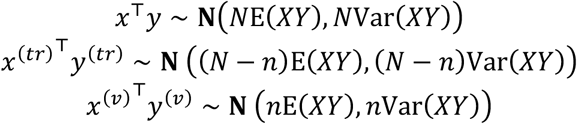

where **N**(*μ*, Σ) denotes a normal distribution with mean *μ* and covariance matrix Σ. It can be shown that

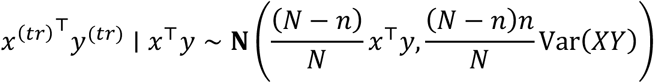

where ⋅ ∣ ⋅ denotes the conditional distribution. This framework does not depend on the linkage equilibrium assumption. However, under simple scenarios where the SNPs are independent (i.e. GWAS summary statistics is pruned), Var(*XY*) is a symmetric matrix whose diagonal and non-diagonal elements can be denoted as

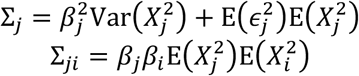

Consequently, we can obtain the validating summary statistics by

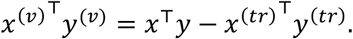

Here, 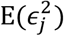 can be estimated by the mean squared error in marginal regressions, which can be further approximated by 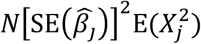. In addition, each SNP’s effect size (i.e. *β*_*j*_) is typically very small in GWAS and 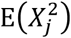 only depends on each SNP’s minor allele frequency (MAF) which is commonly provided in GWAS summary statistics or can be estimated from a reference panel such as the 1000 Genomes Project.^35^ Taken together, Var(*XY*) can be estimated with

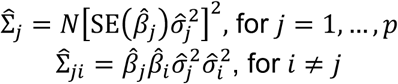

where 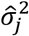 is an MAF-based estimator of 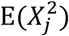. After generating 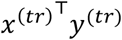 terms as described above from the conditional distribution, the subsampled summary statistics can be estimated by

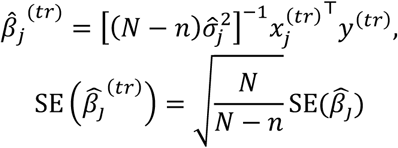

### Evaluating model performance using GWAS summary data

Being able to generate summary statistics for the training and validation datasets resolves a critical issue in model tuning. However, challenges remain in evaluating PRS performance on the testing set without individual-level data. Almost all the PRS approaches in the literature use a linear prediction model as follows

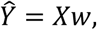

where *w*^T^ = (*w*_1_, …, *w*_*p*_) is the weight for SNPs in PRS. In a traditional PRS, marginal regression coefficients from GWAS are used as the weight values, i.e. 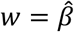, while in other PRS models the weight can be more sophisticated. Here, we demonstrate how to calculate *R*^2^, a commonly used metric to quantify PRS predictive performance, from subsampled GWAS summary data, but our method can be extended to other metrics (e.g. AUC^36^) as well. *R*^2^ on the validation dataset (*y*^(*v*)^, *x*^(*v*)^) can be calculated as

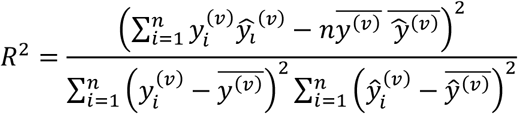

where *ŷ*^(*v*)^ = *x*^(*v*)^*w* and 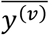 is the sample mean of *ŷ*^(*v*)^. If the SNPs are pruned, it can be shown that the empirical variance of *Ŷ* can be approximated by

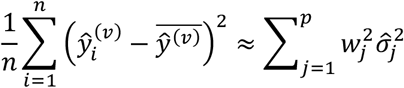

Although empirical variance of *Y* does not affect model tuning, it affects the scale of *R*^2^ and is thus critical for interpreting the results. This term can be approximated by

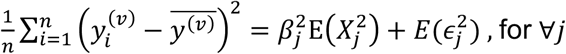

Although Var(*Y*) is always greater than 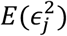 for any *j*, the gap between these two is negligible in real GWAS due to the small effect size of each individual SNP. Thus, a simple estimator for Var(*Y*) can be

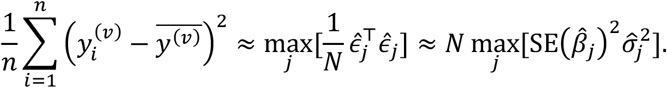

Additionally, since we assumed data to be centered, the mean values in the numerator can be dropped. Taken together, *R*^2^ can be estimated as

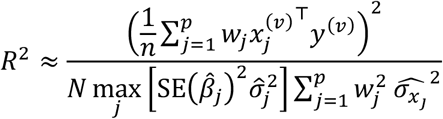

In practice, we use the 90% quantile of 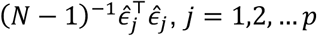, as a more robust estimator for Var(*Y*).

### Model tuning strategies

So far, we have introduced strategies to subsample association statistics on training and validation sets and evaluate model performance using GWAS summary statistics. Combining these two key steps, we will be able perform model tuning using GWAS summary data. Suppose a PRS model uses GWAS marginal estimates 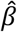 as input and generates SNP weights 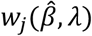 for each SNP. The goal is to find the optimal value of tuning parameter *λ* that maximizes the predictive accuracy. In the simple setting we introduced above, we will generate summary statistics for training and validation datasets. After specifying a tuning parameter *λ*, SNP weights in PRS can be trained by applying the model to the training summary statistics. Then, the prediction accuracy *R*^2^ on the validation summary statistics will be a function of *λ*. Therefore, we can select *λ* so that it maximizes model performance.

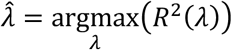

More generally, if the goal is to compare different models, both the summary statistics subsampling and performance evaluation steps remain unchanged. In this case, *R*^2^ will be a function of model and we can choose the best-performing model by optimizing *R*^2^

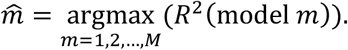

Further, this framework can be used to conduct various types of model tuning procedures. What we have laid out above is the simple training-validation data split approach. If one is interested in applying repeated learning, they can simply repeat the procedure (i.e. resampling training/validation datasets and evaluating *R*^2^ on the validation set) K times. The average *R*^2^ across K folds can be used to select the best model. Similarly, if K-fold cross-validation needs to be implemented, we can first independently simulate K-1 sets of training subsample 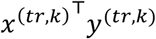 with sample size N/K. Then we can obtain the *K*^*th*^ subsample by

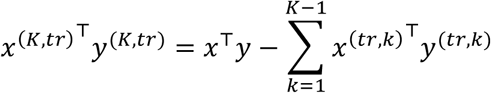

Finally, rotate each one of the K subsamples as a validation sample and the rest as training sample, and use the average *R*^2^ to select the best model. Taken together, PUMAS is a general framework that can perform a variety of model-tuning tasks.

### Simulation settings

We assumed a linear model between a quantitative phenotype and genotypes

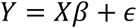

After setting the sample size *N* and the total number of genetic variants *M*, we simulated a *N × M* genotype matrix from independent binomial distribution *Bin*(2, *f*) where *f* is the MAF of each SNP. These simulated genotypes were centered. Then, we generated effect sizes for *m* randomly selected causal variants from a normal distribution **N**(0, (*h*^2^/*m*)*I*), with a predetermined number of causal variants *m* and total heritability *h*^2^. We set the effect sizes of the remaining non-effective SNPs to be 0. Error term *ϵ* were then randomly simulated from **N**(0, (1 − *h*^2^)*I*). Finally, we obtained the values of *Y* by adding up all the components in the linear model.

We performed simulations under a total of 12 different settings. In all settings, we set M to be 5,000 and *f* to be 0.2. Different values were assigned to other parameters, including sample size (*N*=20,000 and 100,000), total heritability (*h*^2^=0.2 and 0.8), and number of causal variants (*m*=50, 1,000 and 4,000). We performed marginal linear regression on the simulated phenotype for each SNP and obtained the summary statistics. We applied PUMAS to perform 4-fold repeated learning on the marginal association statistics. In each fold, 75% of the samples were used as the training set and the remaining 25% were used as the validation set. Average *R*^2^ in four folds of the analysis was used to quantify the model performance. We compared the PUMAS approach to a similar 4-fold repeated learning procedure based on individual-level data – in each fold, we trained PRS on 75% of the samples and calculated *R*^2^ on the rest 25% of samples. We repeated the procedure 4 times and report the average *R*^2^.

### GWAS data

GWAS summary statistics on EA was shared to us by Dr. Aysu Okbay. In this dataset, samples from Add Health, HRS, 23&me, and Wisconsin Longitudinal Study were excluded (N=742,903). Imputed genotype data for Add Health and HRS were accessed through dbGap (phs001367 and phs000428) and the EA phenotypes were defined following the SSGAC GWAS.^37^ 4,775 Add Health samples and 10,214 HRS samples with self-reported European ancestry were used to validate EA PRS. SNPs with imputation quality score < 0.8 were removed from the analysis. The IGAP 2013 AD GWAS dataset was accessed through the IGAP website (http://web.pasteur-lille.fr/en/recherche/u744/igap/igap_download.php). GWAS summary statistics for 7,050 ADGC samples can be accessed through the NIAGADS database (NG00076). Predictive performance on ADGC samples were assessed using summary statistics-based *R*^2^. Following a recent paper, we constructed the AD-proxy phenotype in the UK Biobank based on each sample’s AD status, AD history of parents, whether parents are still alive, and parental age (or age at death).^38^ Imputed genotype data were accessed through the UK Biobank. In addition, we applied PUMAS to benchmark PRS performance on 65 GWASs. Details on these studies are summarized in **Supplementary Table 3**.

For all PUMAS analysis throughout the paper, we first extracted SNPs intersected with the 1000 Genome Phase III data of European ancestry,^35^ Then, we pruned GWAS summary statistics by a LD-block window size of 100 variants, a step size of 5 variants to shift windows, and a pairwise LD (i.e. *R*^2^) threshold of 0.1. We used samples of European ancestry in the 1000 Genome Project Phase III as the reference panel to estimate LD. For GWASs that do not report MAF in the summary statistics, we estimated MAF from 1000 Genome project European samples. In addition, for the analysis of EA and AD, we also intersected GWAS summary statistics with SNPs in the validation set. A p-value grid was used to search for the optimal p-value cutoff (**Supplementary Table 2**).

### Identifying neuroimaging traits associated with AD

GWAS results for imaging traits were accessed from (https://med.sites.unc.edu/bigs2/data/). The IGAP 2019 AD GWAS summary statistics was accessed via NIAGADS (NG00075). We constructed the AD-proxy phenotype in the UK Biobank following a recent paper.^23^ To avoid sample overlap between GWASs, we inferred individuals in the UK Biobank who have undergone brain MRI scans and removed them from the AD-proxy GWAS. All individuals who have visited at least one of the UKB imaging centers were removed from the analysis. 318,773 independent samples remained after removing imaging samples from the data. We performed GWAS with the first 12 principal components,^39^ age, sex, genotyping array, and assessment center as covariates. To test if our approach to remove overlapping samples between neuroimaging GWAS and the AD-proxy analysis was effective, we used cross-trait LD score regression to estimate the intercepts between 211 imaging traits and the AD-proxy GWAS (**Supplementary Figure 4**).^40^ BADGERS software was used to conduct the imaging trait PRS-AD association analysis.^29^ Meta-analysis was conducted using the sample size-weighted approach.^41^

### Code availability

The PUMAS software is available at (https://github.com/qlu-lab/PUMAS).

## Acknowledgements

This project was supported by the Clinical and Translational Science Award (CTSA) program, through the NIH National Center for Advancing Translational Sciences (NCATS), grant UL1TR000427, and by the University of Wisconsin-Madison Office of the Chancellor and the Vice Chancellor for Research and Graduate Education with funding from the Wisconsin Alumni Research Foundation. We thank Dr. Aysu Okbay for generously sharing the EA summary statistics excluding 23andMe, Add Health, HRS, and WLS samples. This study makes use of summary statistics from many GWAS consortia. We thank the investigators for providing publicly accessible GWAS summary statistics. This research uses data from Add Health and HRS. The Add Health Study is supported by Eunice Kennedy Shriver National Institute of Child Health and Human Development Grant P01HD31921 and GWAS Grants R01HD073342 and R01HD060726, with cooperative funding from 23 other federal agencies and foundations. The HRS is supported by National Institute on Aging Grants U01AG009740, RC2AG036495, and RC4AG039029 and is conducted by the University of Michigan. No direct support was received from grant P01-HD31921, R01HD073342, R01HD060726, U01AG009740, RC2AG036495, and RC4AG039029 for this analysis. This research has also been conducted using the UK Biobank Resource under application number 42148. Fletcher acknowledges research support from NIA (P30 AG17266).

## Author contribution

Q.L. conceived and designed the study.

Z.Z., Y.Y., and Q.L. developed the statistical framework.

Z.Z. and Y.Y. performed the statistical analysis.

Y.L. implemented the software.

J.F. assisted in Add Health and HRS data preparation and interpretation.

Y.W. and X.Z. assisted in UK Biobank data processing and analysis.

T.J.H helped interpret neuroimaging risk factors for Alzheimer’s disease.

Q.L. advised on statistical and genetic issues.

Z.Z., Y.Y., and Q.L. wrote the manuscript.

All authors contributed in manuscript editing and approved the manuscript.

