## Supplementary Figures for "Fine-tuning Polygenic Risk Scores with GWAS Summary Statistics"

**Supplementary Figure 1. Comparing the performance of two model-tuning strategies in more simulation settings. (A)** our newly proposed method using summary statistics as input with sample size of 20,000 and heritability of 0.2 **(B)** a traditional repeated learning approach applied to individual-level data with sample size of 20,000 and heritability of 0.2 **(C)** our newly proposed method using summary statistics as input with sample size of 100,000 and heritability of 0.8 **(D)** a traditional repeated learning approach applied to individual-level data with sample size of 100,000 and heritability of 0.8. The X-axis shows the value of tuning parameter (i.e. number of variants in the model). The Y-axis shows the predictive performance quantified by $R^2$. The three curves in each panel represent three different levels of sparsity and genetic architecture.

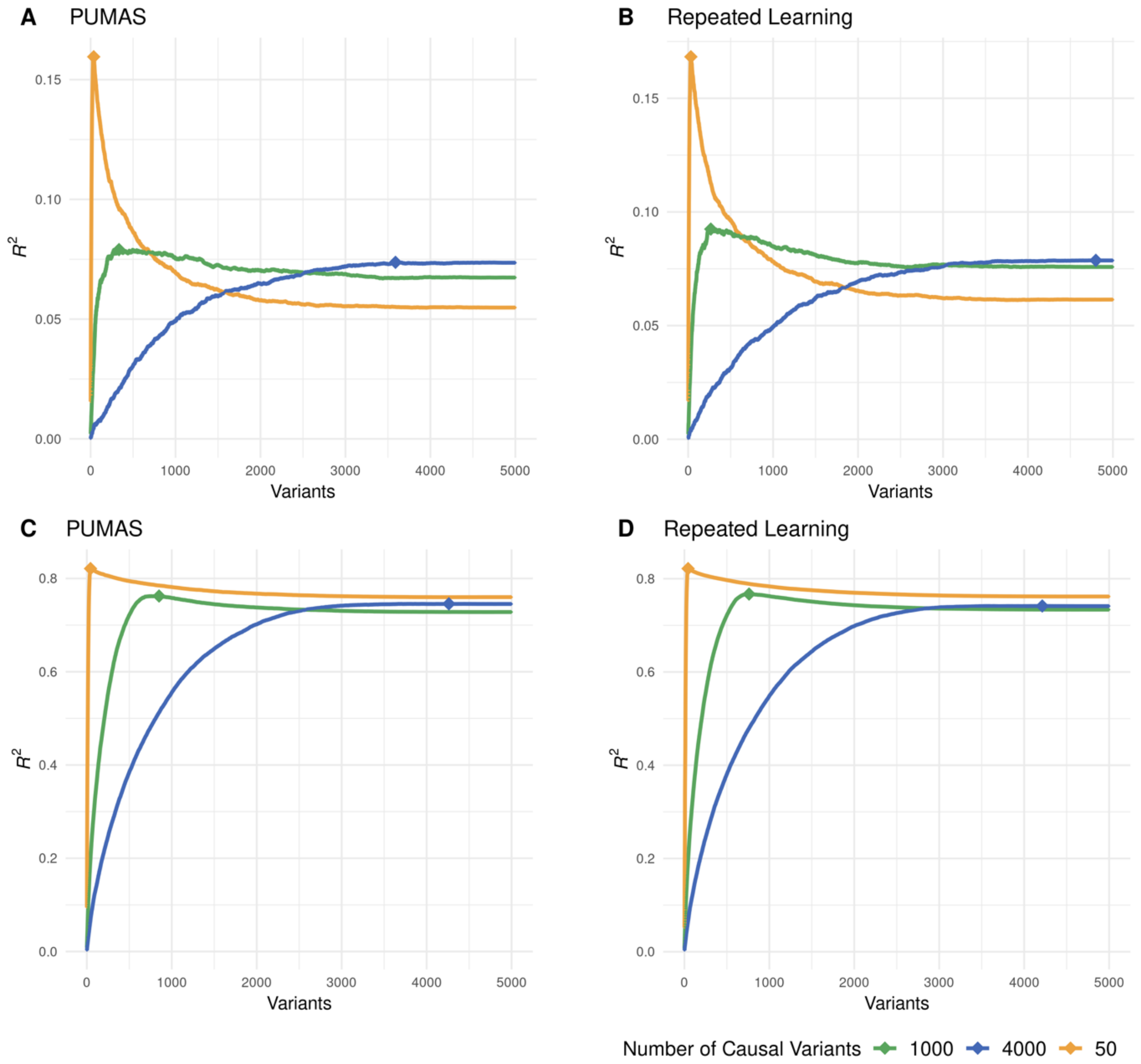

**Supplementary Figure 2. PUMAS result using clumped IGAP 2013 AD GWAS as input.**

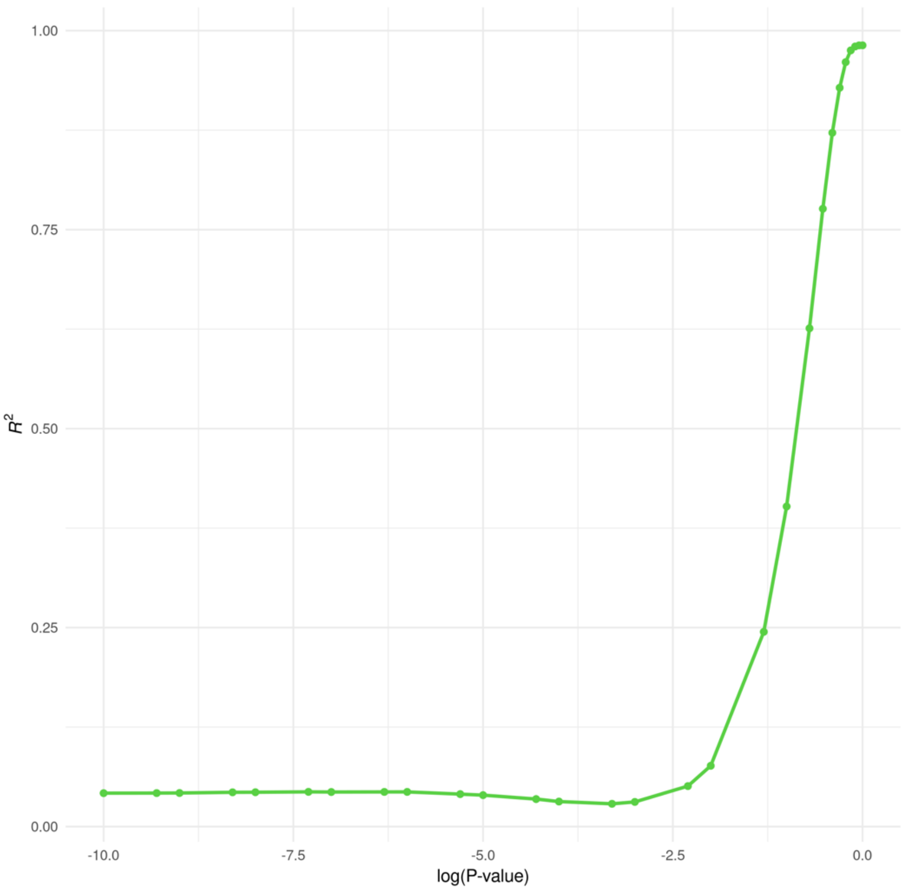

**Supplementary Figure 3. Improvement of predictive $R^2$ of optimized 45 traits. (A)** PUMAS's increase in predictive $R^2$ comparing to PRS of P=0.01 and P=1 **(B)** PUMAS's percentage improvement in predictive $R^2$ comparing to PRS of P=0.01 and P=1. The percentage improvement of RA's predictive performance by PUMAS comparing to its PRS at P=0.01 is truncated to be 2000% in panel B.

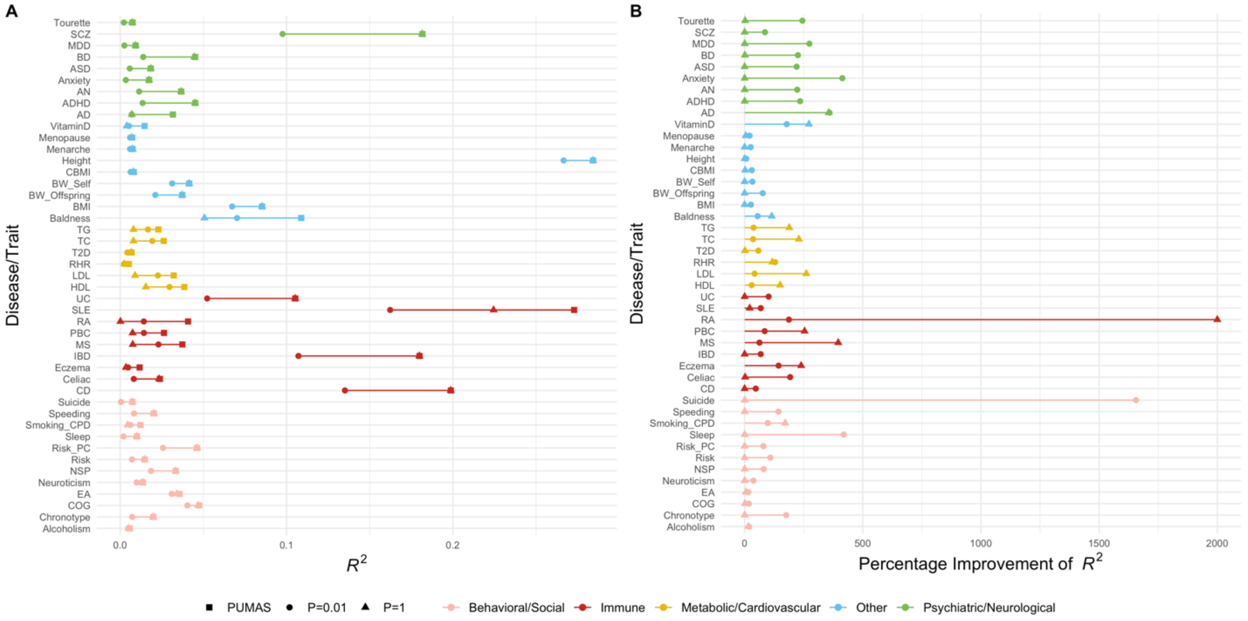

**Supplementary Figure 4. QQ plot for p-values of LDSC intercept estimates between non-imaging AD-proxy GWAS and UK Biobank imaging traits.** P-value for the one-sample t-test of null hypothesis that the mean of LDSC intercepts equals zero is 0.3191.

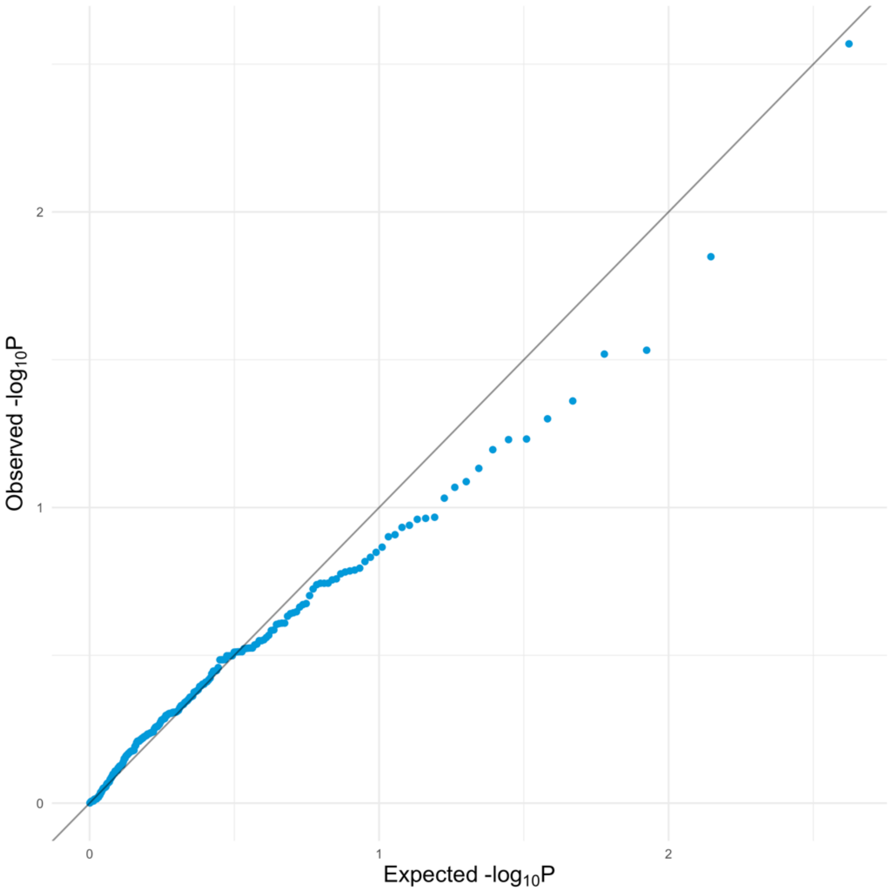

**Supplementary Figure 5. QQ plot for associations between breast cancer and UK Biobank neuroimaging traits.**

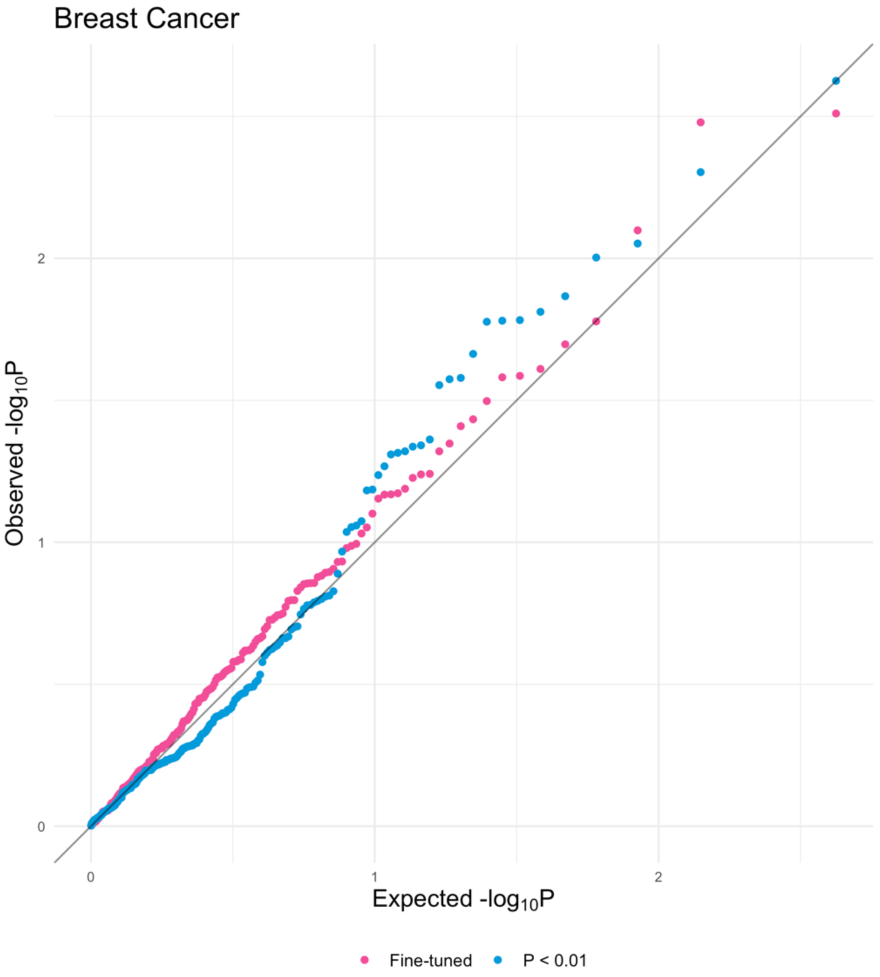
